# Syncytium cell growth increases IK1 contribution in human iPS-cardiomyocytes

**DOI:** 10.1101/862987

**Authors:** Weizhen Li, Emilia Entcheva

## Abstract

Human induced pluripotent stem-cell-derived cardiomyocytes (hiPS-CMs) enable cardiotoxicity testing and personalized medicine. However, their maturity is of concern, including relatively depolarized resting membrane potential and more spontaneous activity compared to adult cardiomyocytes, implicating low or lacking inward-rectifier potassium current (Ik1). Here, protein quantification confirms Ik1 expression in hiPS-CM syncytia, albeit several times lower than in adult heart tissue. We find that hiPS-CM cell culture density influences Ik1 expression and the associated electrophysiology phenotype. All-optical cardiac electrophysiology and pharmacological treatments reveal reduction of spontaneous and irregular activity in denser cultures. Blocking Ik1 with BaCl_2_ increased spontaneous frequency and blunted action potential upstrokes during pacing in a dose-dependent manner only in the highest-density cultures, in line with Ik1’s role in regulating the resting membrane potential. Our results emphasize the importance of syncytial growth of hiPS-CM for more physiologically-relevant phenotype and the power of all-optical electrophysiology to study cardiomyocytes in their multicellular setting.

## Introduction

The growing use of human induced pluripotent stem cell-derived cardiomyocytes (hiPS-CMs) provides genetically-diverse human cardiac models to study cardiac arrhythmia patients with different phenotypes and to assess proarrhythmic risk of new drugs *in vitro* (Savla, Nelson et al. 2014, Shinnawi, Huber et al. 2015). Currently, several cardiomyopathy phenotypes have been successfully recapitulated *in vitro*, including dilated cardiomyopathy (Sun, Yazawa et al. 2012), hypertrophic cardiomyopathy (Lan, Lee et al. 2013) and arrhythmogenic right ventricle cardiomyopathy (Caspi, Huber et al. 2013). Furthermore, these experimental cardiac models are used for discovering new drug therapies (Wen, Wei et al. 2015) and for cardiotoxicity screening (Liang, Lan et al. 2013). This approach overcomes the limitations of species differences in cardiac electrophysiology when using animal models to tackle human diseases (Davis, Casini et al. 2012) and the limitations of studying ion channel macromolecular structures in artificial heterologous expression systems (Hoekstra, Mummery et al. 2012). As the main source of healthy human cardiomyocytes, hiPS-CMs have also been widely used in basic science investigations of human cardiomyocyte calcium handling, metabolism studies, single-cell and multicellular contractility and others (Magdy, Schuldt et al. 2018).

Outstanding concerns about iPS-CMs are related to their level of maturity when compared to adult cardiac tissue. Differences from primary adult cardiomyocytes are seen in cell morphology, calcium handling, glycolysis-based metabolism, and higher cell automaticity (Magdy, Schuldt et al. 2018). While spontaneous contractions of iPS-CM are considered a sign of successful differentiation, high-frequency spontaneous activity is also a proarrhythmic trait in ventricular cells and it influences the iPS-CM’s credibility as a proper *in vitro* model. It has been shown that the spontaneous activity of iPS-CM may be partially influenced by low levels of the inward rectifier potassium current (Ik1) (Vaidyanathan, Markandeya et al. 2016, Li, Kanda et al. 2017). The Ik1 current, encoded by Kir2.1, plays an important role in the stabilization of the cardiac resting potential, the upstroke of the and the late repolarization of the action potential (Figure 1). Insufficient number or low conductance of Kir2.1 ion channels can cause lower membrane resistance (Schanne, Lefloch et al. 1990), with a net inward current during rest, thus a relatively-depolarized resting membrane potential, increased spontaneous rates (Nuss, Marban et al. 1999), slower upstroke and potentially can also affect the action potential duration of hiPS-CMs.

**Figure 1.**
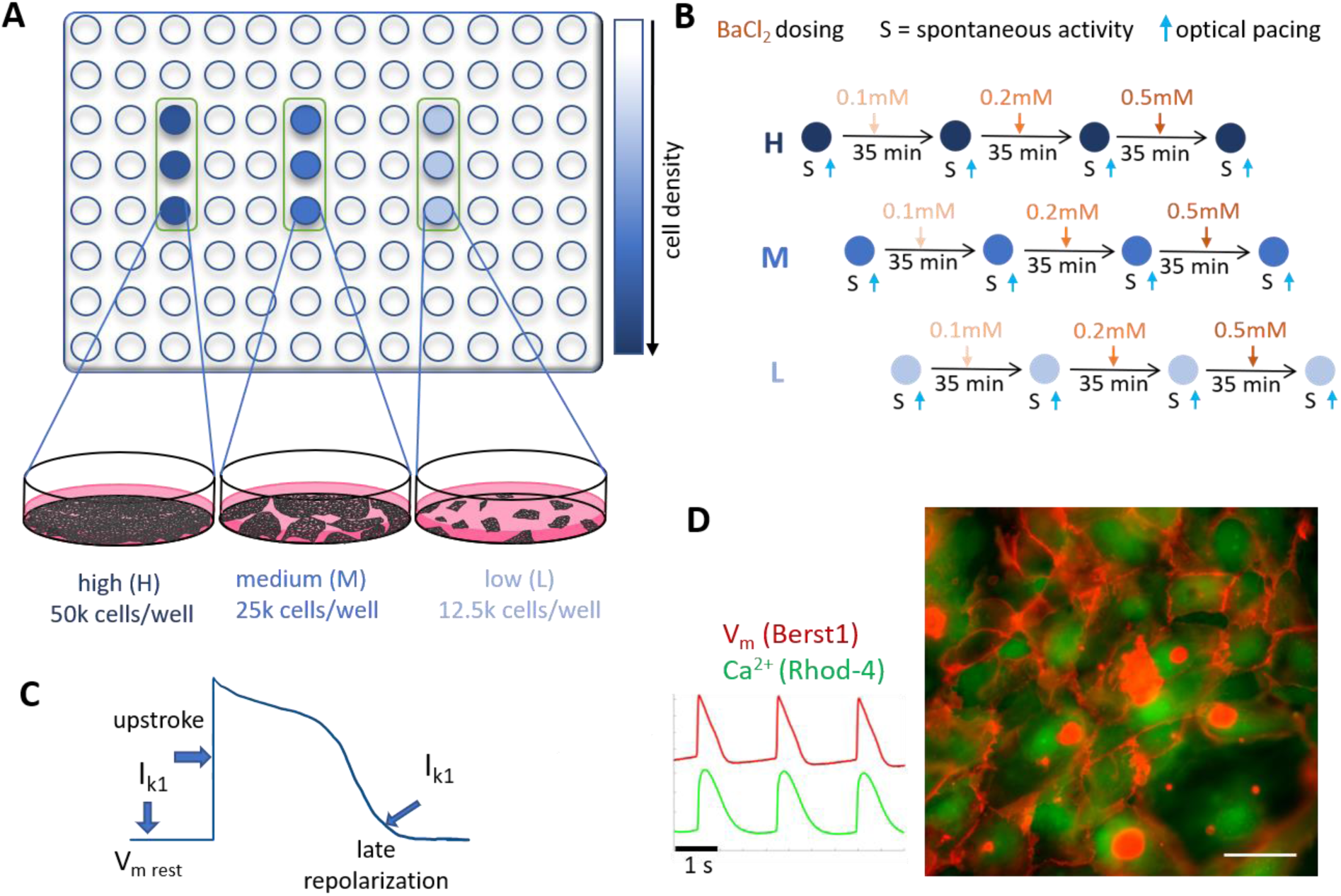
Experimental design using all-optical electrophysiology to probe I_k1_ effects in syncytia of human iPS-CMs. (A) 96-well plate format was used for functional tests. High (H), medium (M) and low (L) cell density were achieved by plating: 50,000, 25,000 and 12,500 cells per well, respectively. (B) Pharmacological probing for I_k1_ contributions by sequential application of increasing doses of BaCl_2_. Functional (voltage and calcium) data were simultaneously collected 35min after each dose treatment under spontaneous conditions (S) and under optical pacing. (C) Expected functional electrophysiological implications of the I_k1_ current: effects on the resting membrane potential, action potential upstroke and contributions to late repolarization. (D) Example image of dual-labeled sample for functional voltage and calcium recordings along with example records. Scale bar is 50μm.

Several approaches have been pursued to increase Ik1 in hiPS-CMs. These include: a). Electrical injection of calculated Ik1-like current via dynamic voltage clamp was applied (Bett, Kaplan et al. 2013, Meijer van Putten, Mengarelli et al. 2015); b). Optical dynamic clamp (ODC) has been applied using light and outward-current-generating opsin ArchT to create a computationally-identical Ik1 contribution (Quach, Krogh-Madsen et al. 2018); and c). Adenoviral overexpression of Kir2.1 has been applied to boost Ik1 in hiPS-CMs (Vaidyanathan, Markandeya et al. 2016, Li, Kanda et al. 2017). Through increased functional contribution of Ik1 (or Ik1-like) current, all three methods claimed a more mature-looking action potential with less depolarized resting membrane potential and suppressed spontaneous activity. As a result, more physiologically-relevant responses to ion channel blockers were reported.

Standard studies of ion channel contributions, including all Ik1 studies in hiPS-CM, are conducted on single cells using voltage-clamp technology. These studies limit the physiological relevance of the obtained functional data, as isolated cardiomyocytes are phenotypically different from cells that exist in a well-coupled cardiac tissue. The cell viability at the end of the cell isolation and the small number of cells that can be studied by such manual techniques inevitably induce selection bias (Hoekstra, Mummery et al. 2012). Previous studies (Du, Hellen et al. 2015, Kane, Du et al. 2016) have drawn attention to the higher variability in cell morphology and function, including action potential morphology, exhibited by hiPS-CMs cultured at low plating density. Other reports have indicated possible effects of cell culture density and cell-cell contacts on gene transcription of specific ion channels (Schanne, Lefloch et al. 1990, Hershman and Levitan 1998, Uesugi, Ojima et al. 2014). These studies prompted us to consider that hiPS-CM multicellular growth conditions may influence cell maturity and the Ik1 contributions to cardiac cell function.

Several voltage-clamp studies of isolated myocytes from adult human heart tissue have reported Ik1 values between 3.6 and 32pA/pF at highly hyperpolarized potentials of −100mV, while values for hiPS-CMs were found to be 10 to 100 times lower (Meijer van Putten, Mengarelli et al. 2015), although recent studies question these findings (Horvath, Lemoine et al. 2018). At these hyperpolarized voltage levels, Ik1 is an inward current and indicates more the power to excite rather than maintain resting membrane potential. The peak outward Ik1 current for isolated myocytes from adult human heart tissue has been reported in the range of 0.4 to 2.2pA/pF (Doss, Di Diego et al. 2012, Meijer van Putten, Mengarelli et al. 2015); the values for the outward Ik1 current in hiPS-CMs have been controversial. Through computations and dynamic clamp studies, it has been shown that peak outward Ik1 influences the resting membrane potential for values under about 3pA/pF, i.e. within the physiological range. To this end, protein quantification and direct comparison between multicellular hiPS-CMs and adult human ventricle has not been done. Furthermore, with a few exceptions, detailed functional interrogations of Ik1’s contributions in multicellular hiPS-CMs are mostly lacking. In this study we investigate the role of cell density on Ik1 and hiPS-CM electrophysiology using protein quantification, all-optical cardiac electrophysiology, pharmacological probing and computational analysis.

## Results

### Syncytia of hiPS-CMs express Kir2.1, albeit at lower levels than adult human tissue; blocking degradation pathways does not rescue Kir2.1 levels

Previous voltage-clamp studies in isolated human iPS-CMs have reported the lack of or insufficient levels of Ik1 compared to adult cardiomyocytes (Meijer van Putten, Mengarelli et al. 2015), with differences at the single-cell level on the order of 10-100 times.

Here, we quantified Kir2.1 protein levels in iPS-CM syncytia using Western Blot and compared those to levels found in various cardiac tissues, including adult human ventricular tissue, adult and neonatal rat ventricular tissues, using wild type HeLa cells as a negative control (Figure 2). Kir2.1 protein was detected at around 48kDa in all samples except wild-type HeLa. Human iPS-CM syncytia were found to have lower normalized protein levels (based on Kir2.1/GAPDH ratio) compared to human heart and rat heart samples. Based on these results (n=4−9 per group), hiPS-CMs’ Kir2.1 expression is about six-fold lower (or about 16.4%) compared to adult human ventricular tissue (Figure 2B). These results confirm previously observed lower Kir2.1 levels in hiPS-CMs, however show less dramatic differences (by an order of magnitude – 6-fold vs. up to 100-fold) compared to results using voltage-clamp in single cells. These findings drew our attention to the possibility that syncytial growth may influence the contribution of Ik1 to electrophysiological function in important ways, and motivated the pursuit of functional experiments in various hiPS-CM growth condition, changing cell culture density.

**Figure 2.**
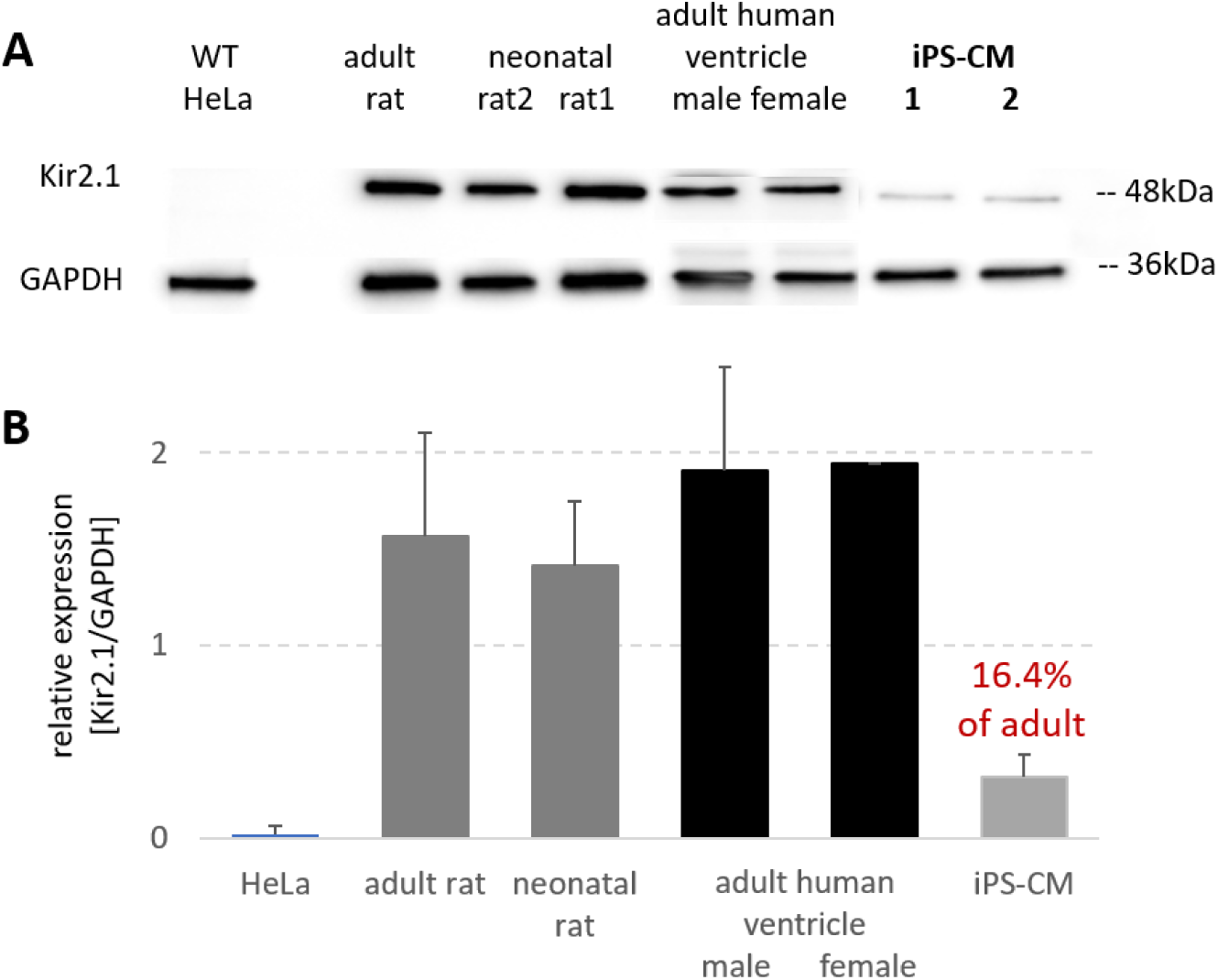
Protein quantification of Kir 2.1 in human iPS-CMs, adult human ventricle, adult and neonatal rat ventricular tissue, and wild-type HeLa cells as a negative control. (A) Western blot results for Kir2.1 protein expression and GAPDH as a loading control. Kir2.1 protein is detected at around 48kDa. (B) Western blot quantification based on Kir2.1/ GAPDH ratio (n=4−9 per sample type, data pooled from five different runs). Tissue samples came from one female and one male human heart, two neonatal and one adult rat hearts; iPS-CM samples came from different cell cultures. Data are normalized by the female human ventricle and shown as mean+/−SD.

To check the possibility that abnormally-fast degradation may be contributing to the lower Kir2.1 protein levels in hiPS-CMs, we applied pharmacological blocking of lysosomal degradation of Kir2.1 (**Suppl. Figure 1**). A previous study (Varkevisser et al. 2013) has demonstrated that dynasore, a non-competitive inhibitor of GTPase activity of dynamin, can inhibit the clathrin-mediated endocytosis of Kir2.1 and preserve it from degradation. We treated syncytial samples of iPS-CMs. Western blot results for 10µM dynasore over 24 hours showed only mild effects on Kir2.1 protein levels (**Suppl. Figure 1**).

### Frequency of spontaneous oscillations in hiPS-CMs increases with lower cell density. Pharmacological probing confirms functional I_k1_ contribution only in denser cell cultures

Functional measurements of voltage and intracellular calcium were carried out in a 96-well plate format using all-optical methods (Figure 1). Three cell density conditions were established by plating 50,000, 25,000 and 12,500 cells per well. Under the high-density condition, well-connected syncytium was formed, with synchronous contractions. Middle-density wells formed less connected cell subgroups with gaps in between. Cells in low cell density wells were more separated, forming smaller groups (**Suppl. Figure 2**), and more likely to exhibit asynchronous subgroup behavior, as seen in other studies as well (Kim, Yang et al. 2015).

Quantification of spontaneous oscillations under different cell plating densities (Figure 3) revealed a decrease in spontaneous beating frequency with the increase of cell culture density, e.g. the low-density group had about 2-fold higher intrinsic frequency of oscillations compared to the high-density cultures. For the sparser cultures, we averaged the frequency of oscillations recorded at multiple clusters within a well.

**Figure 3.**
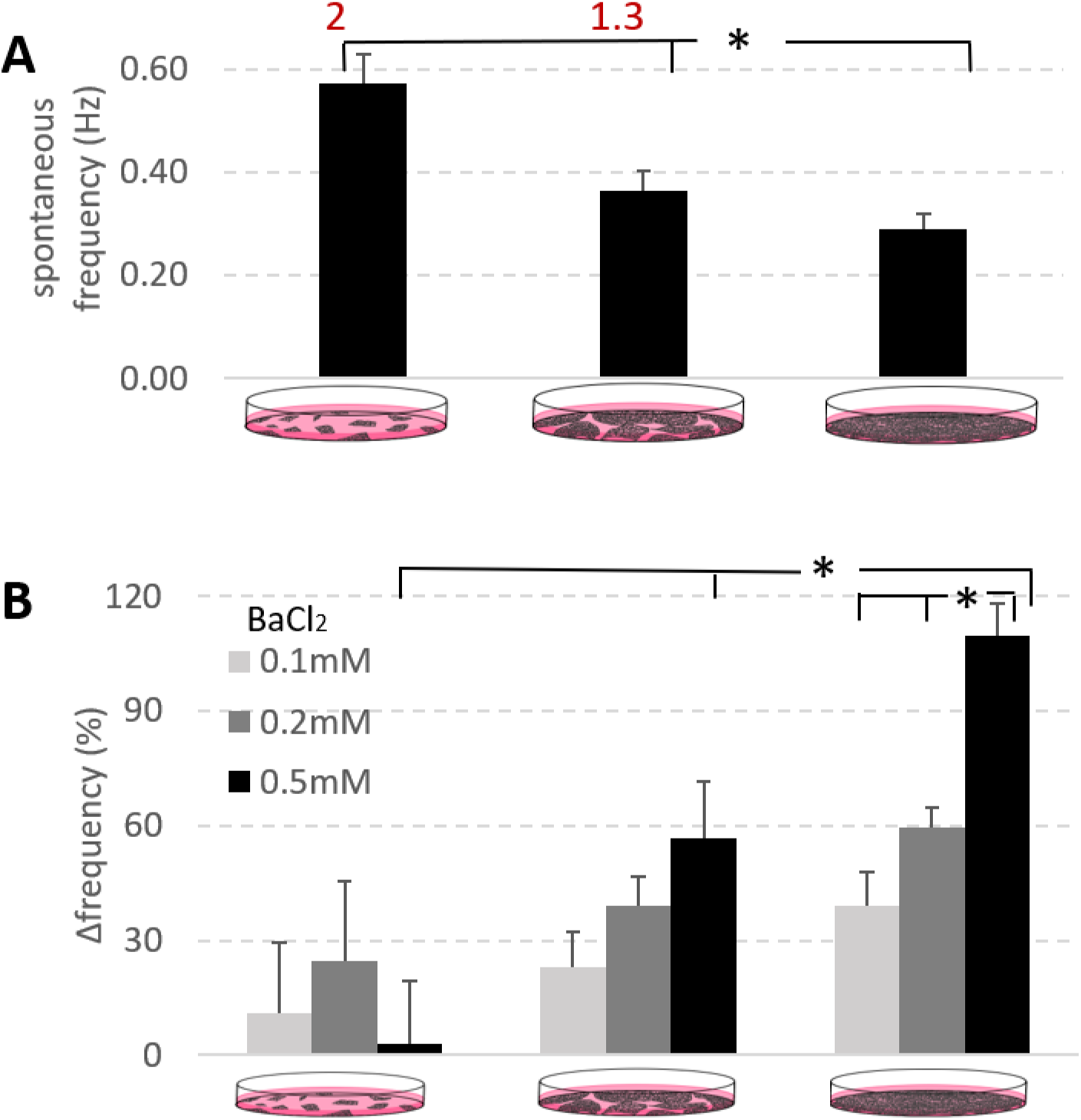
Unmasking I_K1_ contributions to spontaneous activity across different cell culture densities using BaCl_2_. (A) Spontaneous frequency (Hz) of iPS-CM decreases with the increase of cell culture density (*P < 0.05); 2-fold and 1.3 times higher frequency in the low and medium density groups compared to the high-density samples. (B) Changes in spontaneous frequency (%) from baseline are shown for different doses of BaCl_2_. The syncytial high-density group shows a dose-dependent increase in the frequency of oscillations, as expected from I_K1_ block (*P < 0.05). Effects of BaCl_2_ without clear dose dependence in the medium and low-density samples are indicative of minimal I_k1_. Data for this plot were from several locations within n=6 multicellular samples (5 for the low-density group with several isolated regions), subjected to the full sequential drug treatment protocol (12 cases of density and drug dose), data presented as mean+/−SE. Additional experiments with parallel (non-sequential) administration of BaCl_2_ doses were done, yielding similar outcomes, as presented in **Suppl. Fig. 3**.

Considering that low levels of Ik1 are a key driver of spontaneous oscillations in ventricular cells (Schanne, Ruiz-Ceretti et al. 1977, Schanne, Lefloch et al. 1990, Nuss, Marban et al. 1999), we applied a classic pharmacological blocker of Ik1 (BaCl_2_) to the samples with different plating density to test if there will be change in their rates of spontaneous oscillations. The effect on oscillatory activity was quantified as a relative change in spontaneous frequency (%) from untreated conditions, normalized to levels before drug application in the same samples. As summarized in Figure 3B (experiments done according to the sequential dosing described in Figure 1), the syncytial high-density group showed a well-defined dose-dependent increase in the frequency of oscillations, as expected from blocking endogenous I_K1_ (*P < 0.05). In the middle and low-density groups, such dose-dependent changes did not exist or were less pronounced, suggesting little functional contribution of I_k1_. These findings were corroborated in alternative sets of experiments, where BaCl_2_ was applied in parallel to physically-different samples – **Suppl. Figure 3**.

### Lower cell density and suppression of I_K1_ increase occurrences of irregular activity

In addition to the increased frequency of spontaneous activity, the lower-density samples exhibited more irregular activity and variability in the morphology of the action potentials, as seen in other studies as well (Du, Hellen et al. 2015, Kim, Yang et al. 2015, Kane, Du et al. 2016). Irregular activity included: 1) small fluctuations during the resting membrane potential, 2) action potentials with variable amplitude and duration, and 3) embedded lower frequency wave component in the records.

To quantify irregular activity among the different cell density groups, membrane voltage and intracellular calcium were recorded optically. As summarized in Figure 4, no irregular activity was observed in the analyzed traces from the middle and high-density groups under control conditions, while 75% of the low-density cell samples contained such records. Increasing doses of BaCl_2_ to block Ik1 brought about irregular activity in the middle and high-density groups. Again, the high cell density group remained with the least percent of irregular activity events but showed dose dependence. The specific action of BaCl_2_ on Ik1 current and the dose-response in the high-density group potentially support higher functional Kir2.1 expression and contribution compared to lower-density samples.

**Figure 4.**
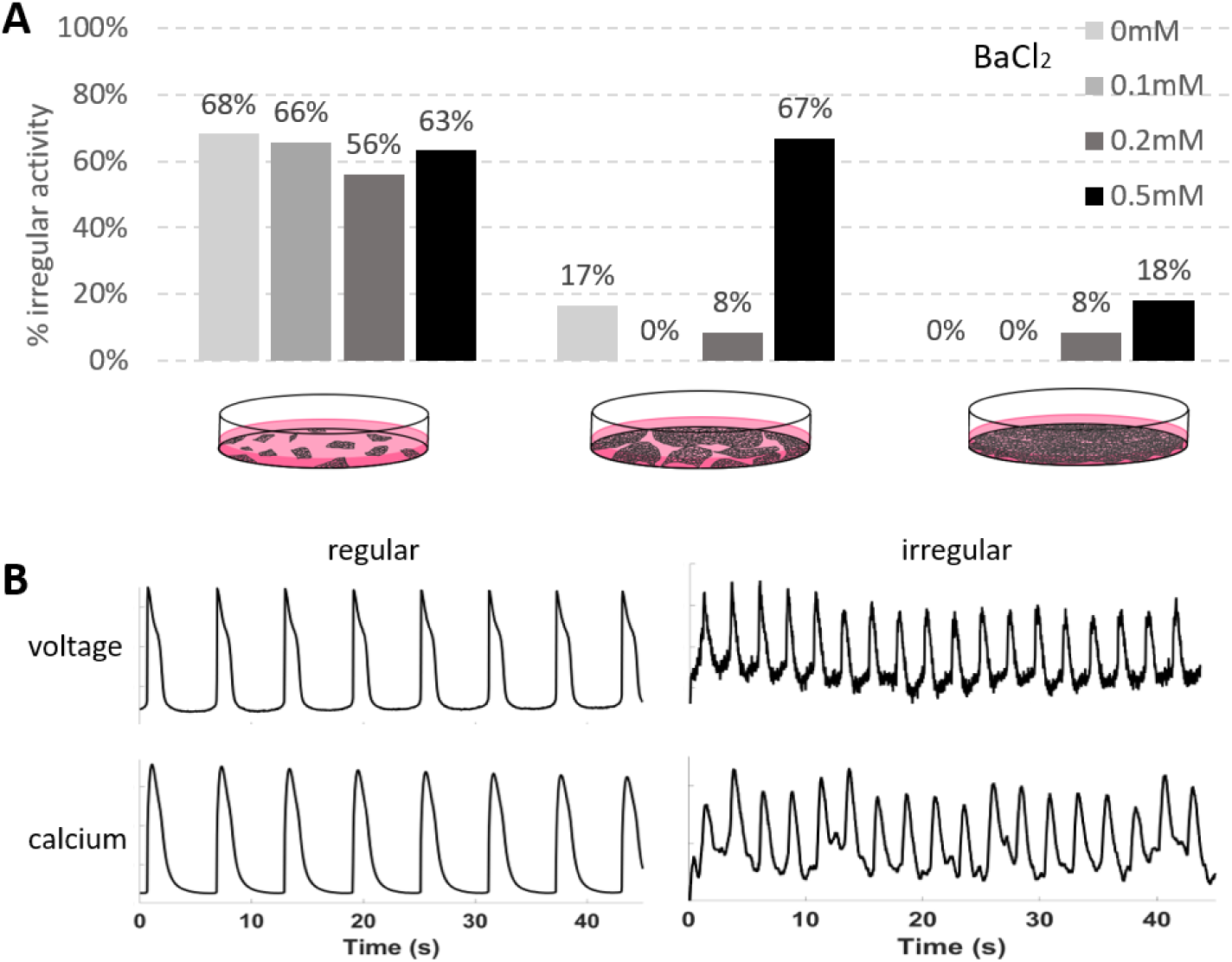
Irregular activity as function of cell density and suppression of I_K1_ by BaCl_2_. (A) Irregular activity is prominent in the low-density group at baseline (no BaCl_2_) and is brought about in the medium and high-density groups by high doses of BaCl_2_. (B) Example traces (spontaneous voltage and calcium) for regular and irregular activity. These results were from multiple records n=11 to 29 per case (12 cases of density and drug dose), see Table of record numbers in the Supplement.

### In paced conditions, I_k1_ levels mainly influence the action potential upstroke, with milder effects on APD

Previous dynamic clamp studies and computational work have indicated effects of Ik1 levels on the morphology of the action potentials in hiPS-CMs (Meijer van Putten, Mengarelli et al. 2015). To study these in controlled conditions, we combined the optical records with optical pacing (Klimas, Ortiz et al. 2019). The recorded paced action potentials (0.7Hz at room temperature) were analyzed by features like action potential upstroke velocity and action potential duration at 90% repolarization (APD90).

Blocking Ik1 with increasing doses of BaCl_2_ in the high-density samples led to a dose-dependent decrease of AP upstroke (Figure 5A), likely due to lower availability of Na+ channels as a result of the more depolarized resting membrane potential. Negligible effects were observed on APD90 in the tested range of Ik1 values (Figure 5B). Computer simulations with a generic adult cardiac ventricular cell model (Luo and Rudy 1994) matched the experimental results considering expected low levels of Ik1 (under 30%, see red-outlined areas), Figure 5C-D. Note that the responses to blocking of Ik1 are different if the cells express higher levels. Similar effects were reported from a dynamic clamp study in hiPS-CMs and computations (Meijer van Putten, Mengarelli et al. 2015) where Ik1 current injection varied from 1 pA/pF to 10 pA/pF. Based on comparison with that study, the red-boxed range of relevant Ik1 values, where AP upstroke is sensitive to Ik1 block and APD90 is not sensitive, likely encompasses peak outward Ik1 values below 3 pA/pF for our syncytial high-density hiPS-CMs; this range includes the values for adult human cardiomyocytes.

**Figure 5.**
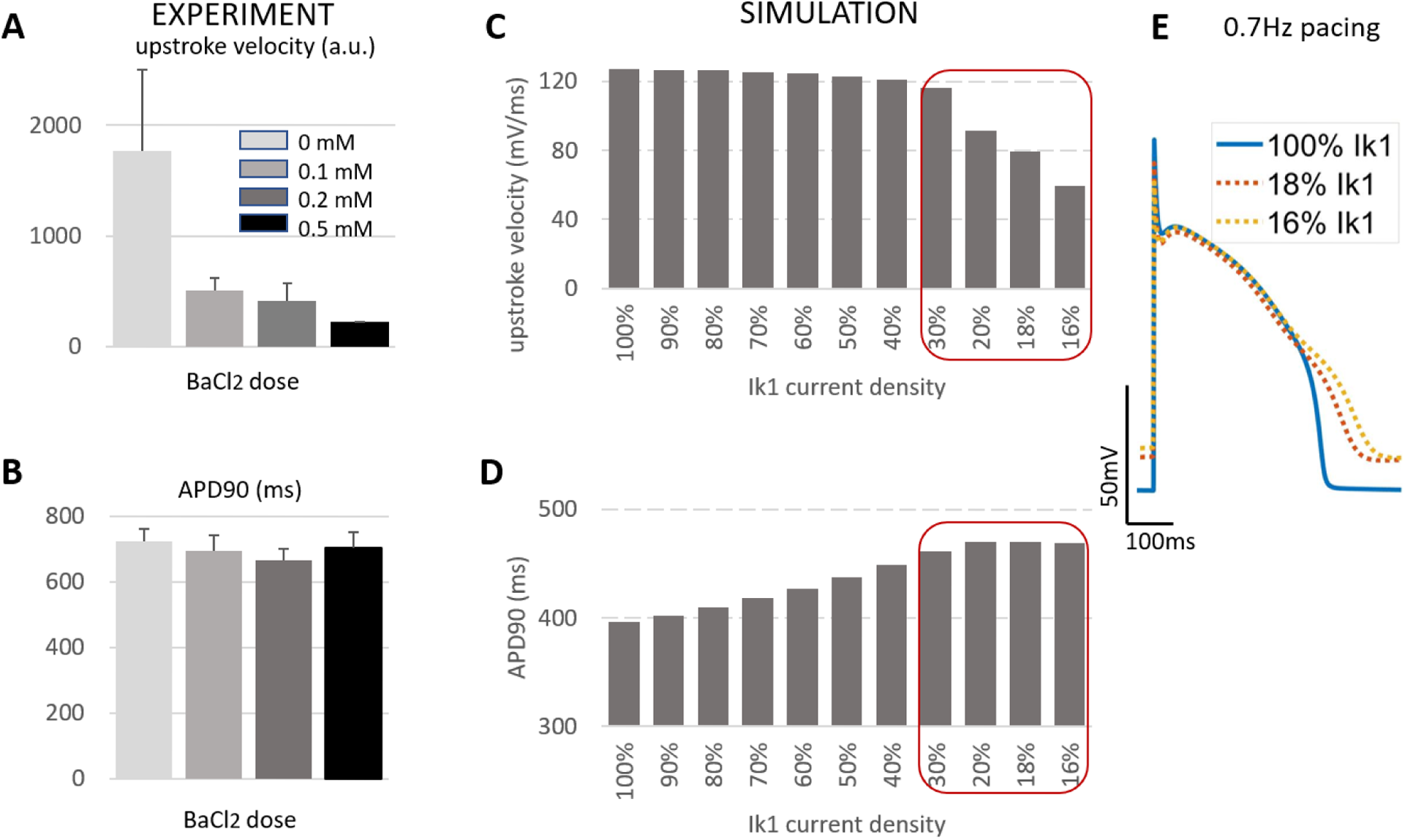
Effects of I_K1_ blocking by BaCl_2_ on optically-paced action potentials in syncytium hiPSC-CMs. Experimentally-obtained (A) maximum upstroke velocity (a.u.) from voltage-sensitive fluorescence dye’s records (∆F/F) of optically-paced action potentials, 0.7Hz; and (B) action potential duration at 90% repolarization (APD_90_) under different BaCl_2_ doses. APD_90_ shows minor changes with a trend towards prolongation with BaCl_2_ treatment. Data are presented as mean+/−SE (n = 4 to 9 high-density multicellular samples for each of the drug doses; multiple locations averaged). Computational results for (C) upstroke velocity in adult ventricular cardiomyocytes as a function of Ik1, analogous to (A); and for (D) APD_90_ in adult ventricular cardiomyocytes as a function of Ik1, analogous to (B). (E) Example traces of simulated action potentials paced at 0.7Hz for different levels of Ik1. Red boxes in (C) and (D) outline the relevant low-Ik1 (<30%) regions of the plots as likely seen in the experimental hiPS-CM data in (A) and (B).

Extending this to variations in cell density and dosing with BaCl_2_, we found that dose-dependent decrease trends for the maximum upstroke velocity exist in the high and middle density groups, but not the low-density group (Figure 6). When comparing the decrease in maximum upstroke velocity before and after 0.5mM BaCl_2_ treatment, we see the biggest difference in the high cell density group, corroborating Ik1 functional contribution in that group.

**Figure 6.**
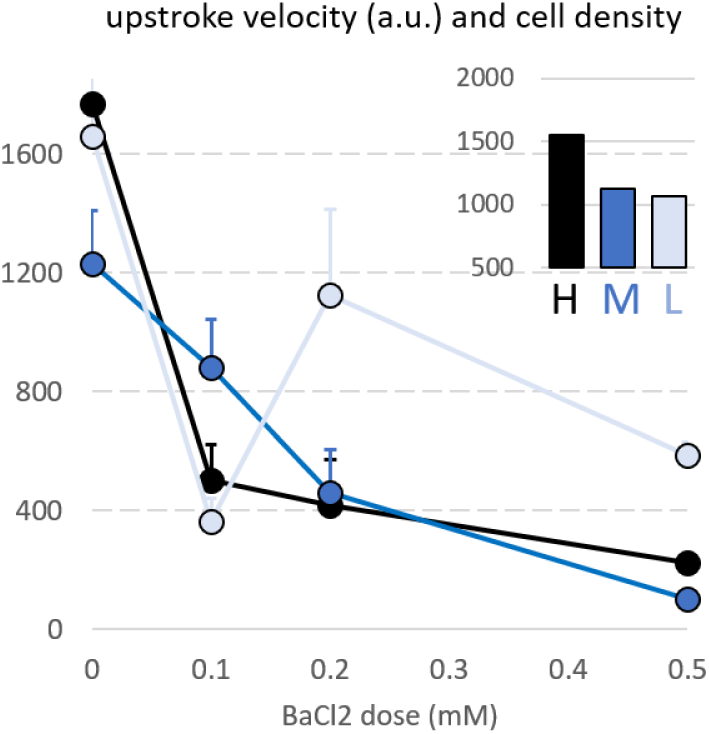
Sensitivity to BaCl_2_ blocking of I_k1_ as function of cell density. Dose response of experimentally-obtained maximum upstroke velocity (a.u.) of optically-paced action potentials when BaCl_2_ is applied. Effects vary between the high, middle and low density iPS-CM cultures. Inset shows the change in upstroke of BaCl_2_ treated (0.5mM) samples and control samples for the three cell densities, with the most pronounced seen in the high-density group. Data (n = 3 to 9 multicellular samples for each of the 12 cases of density and drug dose, multiple locations averaged) are shown as mean+/−SE.

### Cell density also influences responsiveness to blockers of spontaneous calcium release

In addition to the Ik1-mediated mechanism of spontaneous oscillations, activation of the forward mode of the Na^+^/Ca^2+^ exchanger (NCX) and abnormal opening of the ryanodine receptors (RyR) both can increase the probability for spontaneous calcium release (SCR) in cardiomyocytes **(Suppl. Figure 4A)**. If SCR contributes to oscillatory activity along with the low Ik1 levels, blocking the function of either of the calcium-related proteins should decrease cell automaticity. Here, we find that different cell culture densities vary in their sensitivity to the blocking of these channels. As expected, the oscillations in the low-density culture were found more sensitive to blockage of SCR. While 10μM ryanodine caused only about 15% decrease of spontaneous activity in the high-density culture, the low-density group experienced stronger effects. Blockage of NCX by SEA0400 also had higher effect in the low-density culture compared to the high-density culture. The SEA0400 dosage increase from 2μM to 4μM showed relative dose-dependent effects in both groups **(Suppl. Figure 4B)**.

## Discussion

Applications of human iPS-cardiomyocytes for better understanding of human cardiac disease and for personalized medicine require their functional maturation. In the native heart, cardiomyocytes are functioning as well-connected excitable units. Syncytial *in vitro* growth of iPS-CM is expected to help create conditions that are closer to the native environment, compared to isolated cells, and therefore yield a more mature phenotype.

Studies of improving maturity of human iPS-CMs over the last decade span a range of approaches, which include micropatterning, mechanical stretching, electrical field stimulation, induction of cell alignment, genetic manipulations to induce gene expression, delivery of biochemical factors, such as triiodothyronine and alpha-adrenergic agonist phenylephrine or a combination of these (Yang, Pabon et al. 2014). While these strategies of various sophistication levels all may help the maturation of iPS-CMs, the fundamental cell-cell contact effects brought about by syncytial cell growth may represent a simple and impactful way to yield a more mature phenotype for human iPS-CMs.

Here we used protein quantification to establish the expression of Kir2.1 (encoding Ik1) in syncytia of human iPS-CMs, albeit at about 6 times lower levels than in adult human or rat heart tissue (Figure 2). This is in contrast with prior studies in isolated human iPS-CMs that found Ik1 completely lacking (Vaidyanathan, Markandeya et al. 2016, Li, Kanda et al. 2017) or at negligible levels. Further pharmacological investigation (using dynasore as a blocker of clathrin-mediated internalization pathways, **Suppl. Figure 1**) revealed that faster protein degradation is likely not the cause for the lower Ik1 protein levels in hiPS-CMs compared to adult ventricular myocytes, and instead perhaps protein synthesis is different.

To assess the impact of cell density on the contribution of Ik1 to iPS-CM electrophysiology, we created conditions of variable cell density growth and applied all-optical electrophysiology to probe function in these samples. As a fundamental regulator of the resting membrane potential, Ik1 is known to be a key determinant of spontaneous oscillations in ventricular myocytes. Accordingly, we found correlation between the frequency of spontaneous oscillations and the cell density explored in the current study. The highest density syncytial samples showed the lowest frequency of spontaneous oscillations (as mature ventricular cells would) and the lowest occurrence of irregular rhythms (Figures 3 and 4). Furthermore, pharmacological probing of the contributions of Ik1 by variable doses of BaCl_2_ revealed a dose-response only in these high-density samples, while minimal effects were registered in the lower-density samples, with presumably missing Ik1 (Figures 3, 4, 5 and 6, **Suppl. Figure 3**). Comparing experimental data on the effects of lower Ik1 levels on paced action potentials in the hiPS-CMs with computational predictions for adult ventricular myocytes with variable levels of Ik1, we identified key features that are sensitive to Ik1 variations, AP upstroke in particular. Such computations also helped us narrow down the likely range of Ik1 peak outward current in the syncytial human iPS-CM samples to be below 3-4 pA/pF (Figure 6), in line with some previous reports (Meijer van Putten, Mengarelli et al. 2015). Cell density was found to impact also alternative (to the Ik1) potential mechanisms of spontaneous oscillations in the human iPS-CMs, related to spontaneous calcium release, **Suppl. Figure 4**). Frequency of oscillations was more easily suppressed by blockers of NCX and of the RyR receptors in the lower-density samples. Theoretically, in conditions of minimal Ik1 and therefore relatively depolarized resting membrane potential, spontaneous calcium release is expected to more easily lead to voltage oscillations.

Overall, our results draw attention to the importance of syncytial conditions for the electrophysiology of human iPS-CMs; they corroborate previous studies where cell growth density was shown to influence iPS-CM action potential morphology (Du et al. 2015)(Horvath, Lemoine et al. 2018). Furthermore, earlier work with neonatal rat cardiomyocytes demonstrates cell density influence on the expression of potassium ion channels (Guo, Kamiya et al. 1996, Hershman and Levitan 1998). Our results showing that cell density of human iPS-CMs influences Kir2.1 expression and the functional contribution of Ik1 to human iPS-CM electrophysiology is in line with such studies.

A potential mechanism for these effects is the impact that cell-cell contacts have on the expression of ion channels, contributing to differences in the electrophysiology of isolated cells compared to that of cells within a syncytium. Co-localization of ion channels, for example the sodium channel Nav1.5 (encoded by SCN5a), with gap junctional structures (Cx43) has been demonstrated before in cardiac tissue (Kucera, Rohr et al. 2002). Macromolecular complexes of ion channels providing opposing ion fluxes at the cell membrane, e.g. Nav1.5 and Ik1, and their co-localization with the intercalated disks and the perinexus area of cell-cell contacts (Milstein, Musa et al. 2012, Veeraraghavan, Lin et al. 2016) are of particular interest as an efficient mechanism for control of cardiac excitability. Such partnerships and several other known ion channel macromolecular assemblies (Abriel, Rougier et al. 2015) imply that Kir2.1 expression is indeed likely regulated by cell-cell contacts and ion channel clustering, driven by adhesion and gap-junctional proteins, which would only exist in multicellular preparations. This can explain and reconcile our results showing Ik1 contribution in human iPS-CMs only in the denser cell cultures and previous patch-clamp studies reporting lack of Ik1 in isolated cardiomyocytes.

It is important to point out that the classic tools to directly study cardiac electrophysiology, e.g. the patch clamp and voltage clamp, rely primarily on cells in isolation. The limitations of these classic approaches have been pointed out recently (Horvath, Lemoine et al. 2018), specifically in the context of lower seal resistance and ability to assess resting membrane potential when Ik1 levels are low. As in the study by Horvath et al., which analyzed and compared data from isolated cells and from hiPS-CM monolayers and 3D engineered tissues to find similarities between properties of multicellular hiPS-CM samples and adult heart tissue, we highlight the importance of the syncytial setting for cardiomyocytes. Cell dissociation procedures are likely harsher on hiPS-CM aggregates compared to human heart tissue. All-optical cardiac electrophysiology, as used here, offers powerful ways to probe cell phenotypes within the multicellular setting. Albeit not ideal, e.g. in its present form, it cannot report directly contributions of a specific ion current as voltage clamp does, this approach can provide the most comprehensive view of functional responses of hiPS-CMs in a syncytium and can do so in a contactless high-throughput manner (Entcheva 2013, Entcheva and Bub 2016). Further developments may make it possible to also obtain presently missing quantitative ion current information by implementing versions of an optical clamp (Quach, Krogh-Madsen et al. 2018) in conjunction with computational tools.

In summary, this study illustrates that syncytial growth of human iPS-CMs represents a simple approach to influence ion channel expression and obtain a more mature phenotype. All-optical electrophysiology techniques can help track the functional responses of hiPS-CMs and accelerate their application to personalized medicine and optimization for heart regeneration purposes.

## Methods

### Human iPS-CM plating and cell culture

Human iPSC-derived cardiomyocytes (iCell Cardiomyocytes^2^ CMC-100-012-001) from Cellular Dynamics International (CDI), Madison, WI, were thawed based on the manufacturer’s instructions. For protein quantification, cells were plated on fibronectin-coated 6-well plate with recommended plating density of 156,000 cells cm^−2^. For functional recordings, fibronectin-coated 96-well glass-bottom plates were used. Three tested cell culture density are 156,000 cells cm^−2^, 78,000 cells cm^−2^ and 39,000 cells cm^−2^, which represent 50,000 (high density), 25,000 (medium density) or 12,500 (low density) cells per well, respectively. All studies were conducted on days 7-8 after thaw.

### Optogenetic transduction of hiPS-CMs

To enable optical pacing, hiPS-CMs were infected with Ad-CMV-hChR2(H134R)-EYFP 5 days after cell plating, as previously described (Ambrosi and Entcheva 2014, Ambrosi, Boyle et al. 2015). Maintenance medium, containing viral doses of multiplicity of infection (MOI 350), was applied for 2 hours. In the 2 hours period, cells were incubated under 37 °C, 5% CO_2_ and gently agitated every 20 minutes. After 2 hours infection, normal maintenance medium was applied. Functional tests were performed 48 hours after viral delivery.

### iPS-CM protein collection

Protein samples from iPSC-CMs were collected 5 days after cell plating. The whole process was done on ice. After ice cold PBS washes, 120μl RIPA buffer was added in each well and the plate was shaken for 5 minutes. After cells detached from plate, suspended cell content was transferred into microcentrifuge tubes and spun at 15,000 xg for 30 min in a pre-cooled 4°C centrifuge. Clear supernatant was separated for protein quantification.

### Adult human heart tissue processing

Flash-frozen adult human heart tissue was purchased from Amsbio and stored at −80 °C. For protein collection, 30mg of frozen tissue was minced and put on TissueLyser with metal beads and 1ml of RIPA buffer (Thermo Fisher Scientific), run at 50Hz for 2 minutes. Tissue lysis then was spun on 4°C precooled centrifuge for 1 minute to remove the foam, then the TissueLyser step was repeated until tissue was no longer visible; a total of 30min centrifugation at 14,000g, at 4°C.

### Western Blot protein quantification

The total protein amount was quantified using standard BCA assay in all lysates. Then 5-10ug protein per sample was loaded onto 4%-20% gradient gel (Bio-rad) and the gel was run for 2 hours at 90mV. Then bands were transferred onto nitrocellulose membrane using a semidry transfer system (Bio-rad). 5% nonfat milk (Nestle) in TBST buffer (Bio-rad) was used to block the membrane for 1 hour in room temperature. For probing, Kir2.1 antibody (Abcam, ab65796) was diluted 1:200 in 5% nonfat milk in TBST and incubated with the nitrocellulose membrane at 4 degC overnight. The membrane was rinsed with TBST three times, 10 mins each. HRP goat anti-rabbit antibody (Abcam, ab6721) diluted 1:1000 in 5% milk in TBST was applied for 1 hour under room temperature. After three 10 minutes TBST washes, the HRP signal was enhanced by Radiance Plus (Azure) for imaging. Images were taken by Azure C600 Biosystem. After stripping the membrane with striping buffer (15g glycine, 1g SDS, 10 mL Tween20 in 1liter of ultrapure water, adjust pH to 2.2), the membrane was blocked with 5% nonfat milk in TBST again for 1 h in room temperature. GAPDH (Abcam, ab181602) was probed as loading control. 1:1000 GAPDH antibody in 5% milk in TBST was incubated overnight at 4 degC. Same secondary antibody was used to probe the bands and blots were imaged with the same method.

### Functional measurements with all-optical cardiac electrophysiology

Functional experiments were performed using all-optical cardiac electrophysiology as described earlier (Klimas, Ambrosi et al. 2016, Klimas, Ortiz et al. 2019). These were carried out at room temperature in Tyrode’s solution (in mM): NaCl, 135; MgCl_2_, 1; KCl, 5.4; CaCl_2_, 2; NaH_2_PO_4_, 0.33; glucose, 5.1; and HEPES, 5 adjusted to pH 7.4 with NaOH. The optical setup was built around an inverted microscope (Nikon Eclipse Ti2), as described in (Klimas, Ambrosi et al. 2016, Klimas, Ortiz et al. 2019). Optical pacing was done by 470nm light pulses of 5ms, at 0.6-0.8 Hz, using irradiances of 0.4–2 mW mm^2^, as needed. Simultaneous voltage-calcium measurements were performed using temporal multiplexing and iXon Ultra 897 EMCCD camera (Andor Technology Ltd., Belfast, UK), run at 200fps, covering a field of view of about 400μm × 400μm. For spectral compatibility with ChR2, we used a calcium dye Rhod-4AM at 10μM (AAT Bioquest, Sunnyvale, CA) with fluorescence excitation and emission peaks at 530 nm and 605 nm, respectively, and a near-infrared voltage dye BeRST1 at 1μM (from Evan W. Miller, University of California, Berkeley) with fluorescence excitation at 660nm and emission above 680nm, as described in (Klimas, Ortiz et al. 2019). Signals were filtered and analyzed using an automated in-house developed software to extract relevant parameters related to the action potentials and calcium transients under spontaneous and optically-paced conditions (Klimas, Ambrosi et al. 2016, Klimas, Ortiz et al. 2019).

### Blocking Ik1 with barium chloride

Different doses of 0.1mM, 0.2mM and 0.5mM barium chloride (BaCl_2_, Sigma-Aldrich) were applied on each sample serially with 35 min intervals in-between. All-optical electrophysiology was applied to capture spontaneous and optically-paced voltage and calcium from multiple locations after each 35 min treatment. The selected doses of BaCl_2_ for relatively selective suppression of Ik1 were based on prior studies which used 0.1 to 0.5mM (for up to 100% suppression) specifically in human iPS-CMs (Doss, Di Diego et al. 2012, Cordeiro, Nesterenko et al. 2013, Vaidyanathan, Markandeya et al. 2016, Li, Kanda et al. 2017).

### Blocking calcium-dependent pathways with SEA400 and Ryanodine

Suppression of spontaneous calcium release was done by blocking the forward mode of the NCX using SEA0400 (ApexBio, cat no. A3811) at 2μM and at 4μM and by blocking the RyR receptors using Ryanodine (Tocris, cat no. 1329). at 10μM. All-optical electrophysiology was applied to capture spontaneous calcium activity in the low and high-density samples before and 5 min after drug application.

### Blocking clathrin-dependent protein degradation with dynasore

Dynasore (Sigma-Aldrich, cat no. D7693) was dissolved in DMSO at 10mM and stored at −20°C. Upon usage, it was diluted in cell culture media at 10µM applied for 24 hours or 48 hours, based on prior studies (Varkevisser, Houtman et al. 2013). After the treatment, cells were lysed for protein quantification using RIPA buffer.

### Computational modeling of the Ik1 contribution to action potential parameters

Action potential simulations were done using the Luo Rudy’s mammalian ventricular cell mathematical model (Luo and Rudy 1994), which includes the main factors that shape the action potential morphology. In the modeling, the Ik1 current density was varied under S1-S2 stimulation protocol. The measured outputs included action potential upstroke velocity and action potential duration at 90% repolarization (APD_90_).

### Statistical analysis

Comparisons between different groups were done using either single-way or two-way ANOVA with a Tukey-Kramer post-hoc correction for multiple comparisons. Significant differences were considered at p<0.05.

## Acknowledgments

This work was supported in part by an NIH grant R01HL144157 and grants from the National Science Foundation EFMA 1830941 and PFI 1827535 to E.E. We acknowledge help with some experimental aspects by other Entcheva Lab members.

## Author Contributions

WZL performed the experiments, analyzed the data, and produced figures. EE conceived the study and oversaw the project. WZL produced an initial draft of the manuscript, which was edited by EE.

